# Interacting effects of frontal lobe neuroanatomy and working memory capacity to older listeners’ speech recognition in noise

**DOI:** 10.1101/2020.09.14.296343

**Authors:** Nathalie Giroud, Matthias Keller, Martin Meyer

**Author notes:** shared first author (Nathalie Giroud, Matthias Keller). Contact information authors: Dr. Nathalie Giroud Dr. Matthias Keller Prof. Dr. Martin Meyer. **Corresponding author** Dr. Nathalie Giroud, University of Zurich, Department of Computational Linguistics, Phonetics and Speech Sciences, Andreasstrasse 15, CH-8050 Zürich, Switzerland.

## Abstract

Many older adults are struggling with understanding spoken language, particularly when background noise interferes with comprehension. In the present study, we investigated a potential interaction between two well-known factors associated with greater speech-in-noise (SiN) reception thresholds in older adults, namely a) lower working memory capacity and b) age-related structural decline of frontal lobe regions.

In a sample of older adults (N=25) and younger controls (N=13) with normal pure-tone thresholds, SiN reception thresholds and working memory capacity were assessed. Furthermore, T1-weighted structural MR-images were recorded to analyze neuroanatomical traits (i.e., cortical thickness (CT) and cortical surface area (CSA)) of the cortex.

As expected, the older group showed greater SiN reception thresholds compared to the younger group. We also found consistent age-related atrophy (i.e., lower CT) in brain regions associated with SiN recognition namely the superior temporal lobe bilaterally, the right inferior frontal and precentral gyrus, as well as the left superior frontal gyrus. Those older participants with greater atrophy in these brain regions also showed greater SiN reception thresholds. Interestingly, the association between CT in the left superior frontal gyrus and SiN reception thresholds was moderated by individual working memory capacity. Older adults with greater working memory capacity benefitted more strongly from thicker frontal lobe regions when it comes to improve SiN recognition.

Overall, our results fit well into the literature showing that age-related structural decline in auditory- and cognition-related brain areas is associated with greater SiN reception thresholds in older adults. However, we highlight that this association changes as a function of individual working memory capacity. We therefore believe that future interventions to improve SiN recognition in older adults should take into account the role of the frontal lobe as well as individual working memory capacity.

**Highlights:**

- Speech-in-noise (SiN) reception thresholds are significantly increased with higher age, independently of pure-tone hearing loss
- Greater SiN reception thresholds are associated with cortical thinning in several auditory-, linguistic-, and cognitive-related brain areas, irrespective of pure-tone hearing loss
- Greater cortical thinning in the left superior frontal lobe is detrimental for SiN recognition in older, but not younger adults
- Older adults with greater working memory capacity benefit more strongly from structural integrity of left superior frontal lobe for SiN recognition

## 1. Introduction

Many older adults are struggling with understanding spoken language, particularly under adverse listening conditions. For example, older adults have difficulties recognizing speech in the presence of background noise, which makes it hard to participate in spoken conversations and social interactions. For many older adults such hearing problems often mean a reduced quality of life, social isolation, and higher rates of depressive symptoms (Arlinger, 2003; Ciorba et al., 2012; Pichora-Fuller et al., 2015; Vannson et al., 2015). Furthermore, hearing impairment in older adults is associated with cognitive decline (de la Fuente et al., 2019; Fischer et al., 2016; Fortunato et al., 2016; Merten et al., 2019) as well as a higher risk for incident all-cause dementia (Albers et al., 2015; Deal et al., 2019, 2015; Gates et al., 2011; Lin et al., 2011; Lin and Albert, 2014; Osler et al., 2019).

Speech-in-noise (SiN) recognition difficulties in older adults can partly be explained by the highly prevalent pure-tone hearing loss (i.e., elevated pure-tone thresholds, particularly in higher frequencies) (Cruickshanks et al., 1998; Dubno et al., 1984; Gordon-Salant and Fitzgibbons, 1999; Killion and Niquette, 2000). Age-related pure-tone hearing loss often results from a damage to cochlear outer hair cells and the stria vascularis in the auditory periphery (Dubno et al., 2013; Mills et al., 2006). But also older adults without elevated hearing thresholds, or when audibility is restored with amplification, have considerably lower tolerance for background noise during speech recognition compared to younger adults (Füllgrabe, 2013; Füllgrabe et al., 2015; Giroud et al., 2018; Hopkins and Moore, 2011; Moore et al., 2014; Pichora-Fuller and Souza, 2003). Thus, other factors, apart from pure-tone hearing loss, may account for variance in SiN reception thresholds in older adults.

In an important position paper by Humes and colleagues (2012) it was proposed that, in addition to pure-tone hearing loss, age-related changes in the (sub)cortical auditory-related brain areas as well as age-related decline in cognitive capacity, may contribute to the observed decline in SiN recognition in older adults and hence to impairments in every day conversation. Evidence from structural neuroimaging studies indicated that age-related atrophy in the right Heschl’s gyrus (HG) contribute to SiN recognition problems in older adults (Giroud et al., 2018). Decline in brain structure (i.e., cortical thinning and loss of cortical volume) across the lifespan is part of a normal aging process, while considerable interindividual variability has been described (Fjell et al., 2014, 2009; Storsve et al., 2014; Walhovd et al., 2011). Age-related atrophy in primary auditory but also in non-primary auditory areas, as well as in brain regions involved in language processing and cognition, have been associated with greater SiN reception thresholds (Bilodeau-Mercure et al., 2015; Giroud et al., 2018, in prep.; Rudner et al., 2019; Tuwaig et al., 2017; Wong et al., 2010) and other speech perception tasks (Giroud et al., 2019). For example, lower cortical volume and reduced thickness of the left and right superior temporal gyrus (STG), of Heschl’s sulcus (HS) (Giroud et al., 2018; Rudner et al., 2019), of the inferior frontal gyrus (IFG) (i.e., of the left pars orbitalis and of the left pars triangularis), and of the left prefrontal cortex (PFC) (i.e., left superior frontal gyrus) (Giroud et al., 2018; Wong et al., 2010)) have been reported to be related to greater SiN reception thresholds. While the STG, the HS, and the IFG are generally considered part of the core language network (for overviews, see e.g. Friederici, 2012; Hickok and Poeppel, 2007, 2004), superior prefrontal regions are usually considered to be involved in at least some principal cognitive functions such as executive control and working memory (Elliott, 2003).

Functional neuroimaging studies have revealed that a stronger involvement of the PFC during SiN recognition might be related to cognitive compensatory mechanisms. A study by Wong and colleagues (2009) found increased BOLD-related neural responses in the PFC and the precuneus during SiN processing in older compared to younger adults with normal pure-tone hearing. The authors interpreted this neural pattern to reflect greater utilization of the phonological working memory in older adults reflecting the increased speech processing demand due to the background noise. A meta-analysis of auditory neuroimaging studies further supports this idea showing that higher speech processing demand due to background noise is associated with greater brain responses in the left IFG, among other brain regions (Alain et al., 2018). It was hypothesized that, when the STG is not able to process speech sounds effectively due to the presence of background noise, cognitive functions such as executive control, inhibition, attention, working memory and speech-motor integration in the PFC may be recruited to eventually achieve an interpretation of the spoken utterance (Du et al., 2016; Rudner et al., 2019).

Similarly, the Ease of Language Perception (ELU) model (Rönnberg et al., 2019, 2013, 2008) highlights the importance of cognition during SiN perception. Based on research showing that higher working memory capacity is associated with improved speech recognition, particularly in adverse listening situations (Anderson et al., 2013; Dryden et al., 2017; Wingfield et al., 1998), the ELU model describes how working memory and particularly the episodic buffer become important in speech perception when there is a mismatch between the auditory input and the stored representation of sounds in the semantic long-term memory. Such a mismatch can occur, for example, due to background noise or hearing loss. When distorted or unclear auditory input does not match with phonological or lexical representations, working memory functions keep the relevant auditory input active, while more information such as contextual cues can be processed until a match between the input and the stored representations can be achieved. Thus, age-related decline in working memory capacity may lead to lower SiN recognition abilities in older adults.

In sum, there are two separate lines of research evidencing, that - apart from pure-tone hearing loss - 1) age-related atrophy in auditory- and cognitive-related brain regions and 2) age-related decline in working memory may lead to greater difficulty with SiN recognition in older adults. In the present work, we combined these two separate, but not independent, lines of research. We analyzed to what degree age-related brain atrophy, especially in prefrontal areas, predicts SiN performance in a group of healthy older adults with age-appropriate pure-tone hearing and compared their measurements with those obtained from younger controls. Furthermore, we assessed the older adults’ working memory capacity in order to estimate its role in the association between atrophy and SiN performance using a moderation analysis. We hypothesized that older adults have greater atrophy in auditory (e.g. HG, STG, STS) and cognitive-related brain regions (e.g. PFC) reflecting lower SiN performance as compared to younger adults. Furthermore, we predicted a moderating effect of working memory capacity on the association between prefrontal atrophy and SiN performance in older adults. We believe that the investigation of a potential moderation of individual working memory abilities will help to better understand the role of prefrontal areas for speech perception in older adults.

## 2. Methods and Materials

### 2.1. Participants

Twenty-five older adults (OA) (age range = 65-80 years, M_age_=70.88, 12 females) and 13 younger adult (YA) controls (age range = 20-29 years, M_age_=24.15, 10 females) participated in this study. All participants scored at least 27 points in the Mini-Mental State Examination (MMSE) (Folstein et al., 1975) ensuring that participants did not suffer from dementia. Furthermore, the volunteers reported to not have any history of psychological or psychiatric disorders or brain injuries. None of the participants were using hearing aids and they all reported to not have any serious speech or hearing impairments. Both, YA and OA, were native (Swiss-) German speakers and did not learn any second language before the age of seven years. The Annett Hand Preference Questionnaire (Annett, 1970) indicated that all participants were right-handed. Participants were paid for their participation and gave informed written consent. Data collection has been approved by the ethics committee of the Canton of Zurich.

### 2.2. Pure-tone thresholds

Audiometric testing was performed in a double-walled, sound-attenuated booth at the University Hospital of Zurich. Pure-tone thresholds were measured for 500, 1000, 2000 and 4000 Hz using a probe-detection paradigm in which pure tones were presented for 250 ms (Lecluyse et al., 2013; Lecluyse and Meddis, 2009). Tones were delivered via a custom-written Matlab software to circumaural headphones (Sennheiser HD 280-13 300Ω). Participants’ responses were recorded with a touch screen (ELO AccuTouch, version 5.5.3.6.). Only participants with pure-tone thresholds below 40 dB were included in this study in order to rule out unwanted effects of moderate or severe pure-tone hearing loss. While the YA did not show any hearing impairment according to the hearing loss categorization of the World Health Organization, the OA had elevated thresholds that ranged between none (<= 25 dB) to mild impairment (<= 40 dB) (see Figure 1).

**Figure 1:**
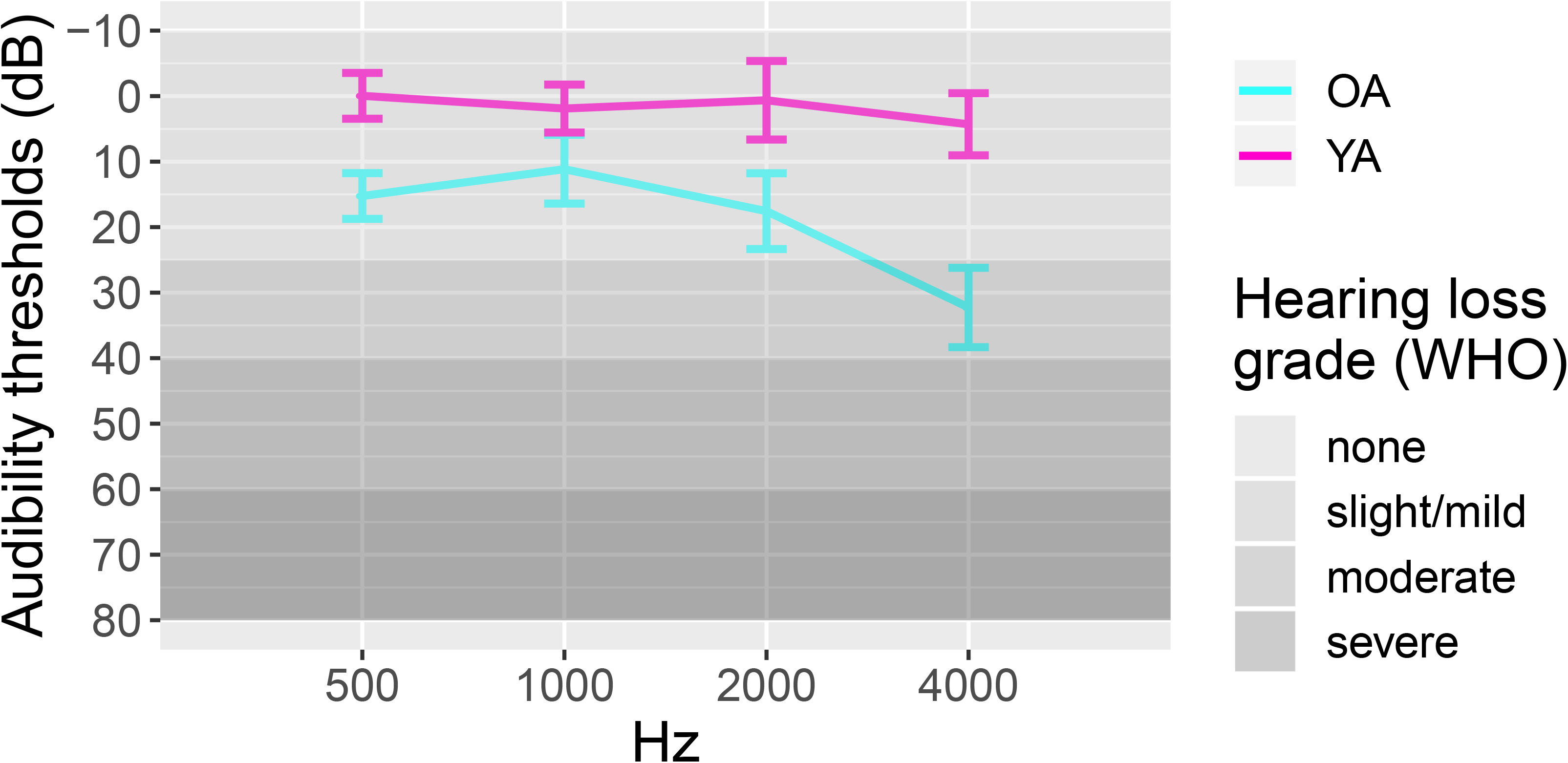
Pure tone hearing thresholds averaged over both ears of the young and the older age groups. The World Health Organization (WHO) classification of hearing loss is displayed on the right. Error bars indicate 95% confidence intervals.

### 2.3. Speech-in-noise recognition

SiN recognition was assessed by means of the OLSA Matrix Sentence Test (Wagener et al., 1999c, 1999a, 1999b). In the OLSA, SiN reception thresholds are computed with an adaptive approach. Sentences and noise were presented simultaneously to the participants. After each sentence, participants were asked to repeat as many words as possible from the preceding sentence. Sentences were low-context sentences to prevent participants from guessing the correct answer due to the sentences’ context. The noise was generated by 30 overlays of the whole test material, which led to low amplitude modulation noise in the same spectrum as the test sentences. At the beginning of the testing procedure, sentences and noise were presented at 65 dB SPL. After that the sentence level was varied until the participant was able to correctly repeat 50% of the words of the sentence. Participants were seated in front of a loudspeaker which was positioned 0° azimuth and 1.5 m away from the person’s head. Sentences were presented via in-house developed MACarena software (https://www.uzh.ch/orl/projects/speechtests/speechtests.html).

### 2.4. Working memory

Working memory performance was assessed in the older group with a verbal auditory *n*-back paradigm (Nystrom et al., 2000; Owen et al., 2005). Letter sounds were presented from a computer and participants were asked to identify whether the letter was the same as the one from n steps earlier. Each participant completed a 2-back and a 3-back run, each consisting of 50 letters including 19 matches in total. Inter-stimulus-interval between letters was four seconds and participants indicated a match to the letter presented n steps back by pressing the space bar. Participants performance was defined as the ratio between total responses given and correct responses for both runs.

### 2.5. MR acquisition and image processing

Structural MR images were recorded using a T1-weighted turbo field echo sequence (160 sagittal slices, in-plane resolution = 0.94 × 0.94, slice thickness = 1 mm, matrix size = 256 × 218 × 256 mm, FOV = 240 × 240 mm, repetition time [TR] = 8.15 ms, TE = 3.74 ms, flip angle = 90°). T1-weighted images were analyzed with the FreeSurfer image analysis suite (version 6.0.0., http://freesurfer.net/). Using the surface-based morphometry (SBM) approach implemented therein, cortical surface models of all participants were obtained automatically (Dale et al., 1999; Fischl et al., 2004, 2002, 2001, 1999a, 1999b; Ségonne et al., 2004). After segmentation, surface reconstructions were checked for accuracy. For one subject, white matter segmentation errors were found, which were then corrected manually. The resulting surface models yielded measures of cortical thickness (CT) and cortical surface area (CSA). Hereby, CT denotes the shortest distance between the gray/white matter border and pial surfaces and CSA the mean area of the triangular region at the respective vertex. Each participant’s reconstructed brain was morphed to an average surface and smoothed using a FWHM kernel of 10 mm (Meyer et al., 2016). These were then entered to the built-in general linear model (GLM) facility of FreeSurfer for statistical analyses.

### 2.6. Statistical analyses

Statistical analyses were performed using the R software (R Core Team, 2017), version 3.4.2. First, it was tested to what degree age, PTA, and working memory were associated with SiN performance. Participant’s age and PTA were entered as predictors into a multiple regression model with the SiN threshold as a dependent variable, while controlling for sex. Working memory (i.e., n-back hits) and SiN were correlated, controlling for PTA and sex. Second, brain regions for which CT or CSA were associated with SiN performance were identified across all participants by entering the SiN threshold as a regressor in the FreeSurfer GLM facility with the whole-brain anatomical measures as dependent variables. The resulting models were corrected for multiple comparison by applying Monte Carlo Null-Z simulation with a vertex-wise/cluster-forming threshold of *p*<.001 and a cluster threshold of *p*<.05 for each hemisphere independently. Anatomical measures of the significant clusters were then extracted and subjected to further analysis as follows: 1) Independent samples *t* tests were performed to compare CT and CSA measurements between the two age groups for each anatomical region. In this case *p*-values were Bonferroni corrected for the number of regions compared within each anatomical measure. 2) To assess in which brain regions age moderated the relationship between anatomical measures and SiN performance, multiple linear regression models were calculated with SiN performance as dependent variable and anatomical measures as predictors and age as a moderator, while sex and PTA were included as covariates. 3) To assess to what extent working memory moderated the association between neuroanatomical traits and SiN in older adults, the interactions between SiN performance and z-standardized n-back performance were entered as predictors with the anatomical region of interest as dependent variable into a multiple linear regression model with sex and PTA as covariates.

## 3. Results

### 3.1. The association between age and SiN performance

Age (*β*=0.85, *p*=.039) significantly predicted SiN performance (F(3,34)=13.33, *p*<.001, adjusted R^2^=0.5), suggesting that older individuals’ ability to process speech masked by background noise was worse than in younger adults (see Figure 2). PTA did not significantly predict SiN recognition in our sample. Our results therefore show that SiN performance drops significantly with age, irrespective of pure-tone hearing loss (SiN performance (dB SNR) in younger controls: M=−5.66, SD=.56, range=−6.7−.5.0; older adults: M=−2.42, SD=1.70, range=−7.0−1.8). SiN performance did not significantly correlate with our n-back working memory measure (*r* =.13, *p* =.55) in our sample of older adults (n-back hits in older group: M=2.65, SD=1.20, range=1.22−6.50). In the next sections, we assessed to what degree age-related atrophy was related to the lower SiN performance only in older adults.

**Figure 2:**
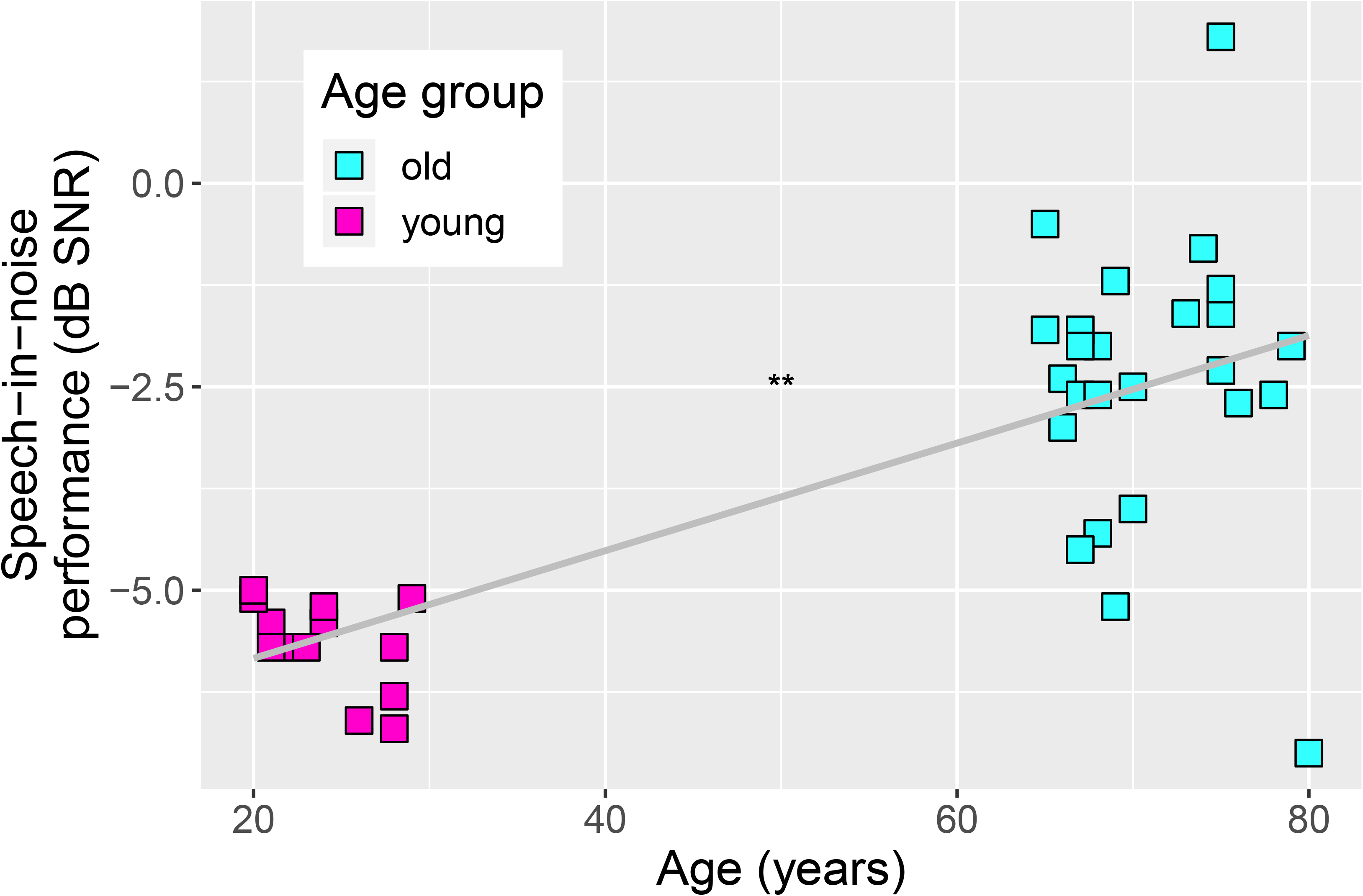
Significant positive association between age and speech-in-noise reception thresholds showing that older adults perform worse in SiN than younger adults. \*\**p*<.01.

### 3.2. Associations between cortical thickness and cortical surface area and SiN performance

In order to test to what degree cortical surface area and cortical thickness were related to SiN performance, a whole-brain FreeSurfer GLM analysis was computed. It revealed several clusters in the left and right hemisphere for which CT was significantly correlated with SiN performance irrespective of age, while no associations were found for CSA. The cluster statistics for CT are presented in Table 1 and Figure 3 with the annotations drawn from FreeSurfer. Significant associations between CT and SiN were found in superior temporal regions bilaterally, the right inferior frontal cortex, the right caudal middle frontal cortex and right precentral regions. In the left hemisphere, significant correlations were found with the superior frontal lobe. CT in all clusters was negatively correlated with SiN indicating that individuals with thicker cortices in these regions performed better in the SiN task.

**Table 1:** Significant correlations between cortical thickness (CT) and speech-in-noise (SiN) reception thresholds in the two hemispheres irrespective of age group. All correlations were negative showing that those with greater CT performed better in SiN. LH: Left hemisphere, RH: Right hemisphere, Max: log10(p) at peak vertex (values < −3 correspond to p < 0.001), NVts: number of vertices above threshold (*p* < 0.001, cluster-corrected).

|  |  |  |  | Talairach coordinates |  |  |
| --- | --- | --- | --- | --- | --- | --- |
| Annotation | Max | NVtxs | Size (mm2) | X | Y | Z |
| LH |  |  |  |  |  |  |
| Superior frontal | -5.079 | 1234 | 676.34 | -7.6 | 56.8 | 17.7 |
| Superior temporal | -5.804 | 930 | 428.28 | -44.5 | -9.5 | -13.0 |
| RH |  |  |  |  |  |  |
| Superior temporal | -4.468 | 722 | 302.81 | 41.9 | -12.8 | -10.3 |
| Pars triangularis | -5.449 | 488 | 295.78 | 48.4 | 34.2 | -1.8 |
| Superior temporal | -4.911 | 616 | 259.52 | 42.3 | -34.0 | 14.2 |
| Caudal middle frontal | -4.522 | 380 | 251.51 | 35.2 | 18.9 | 44.1 |
| precentral | -6.614 | 474 | 223.33 | 56.5 | 5.8 | 11.5 |

**Figure 3:**
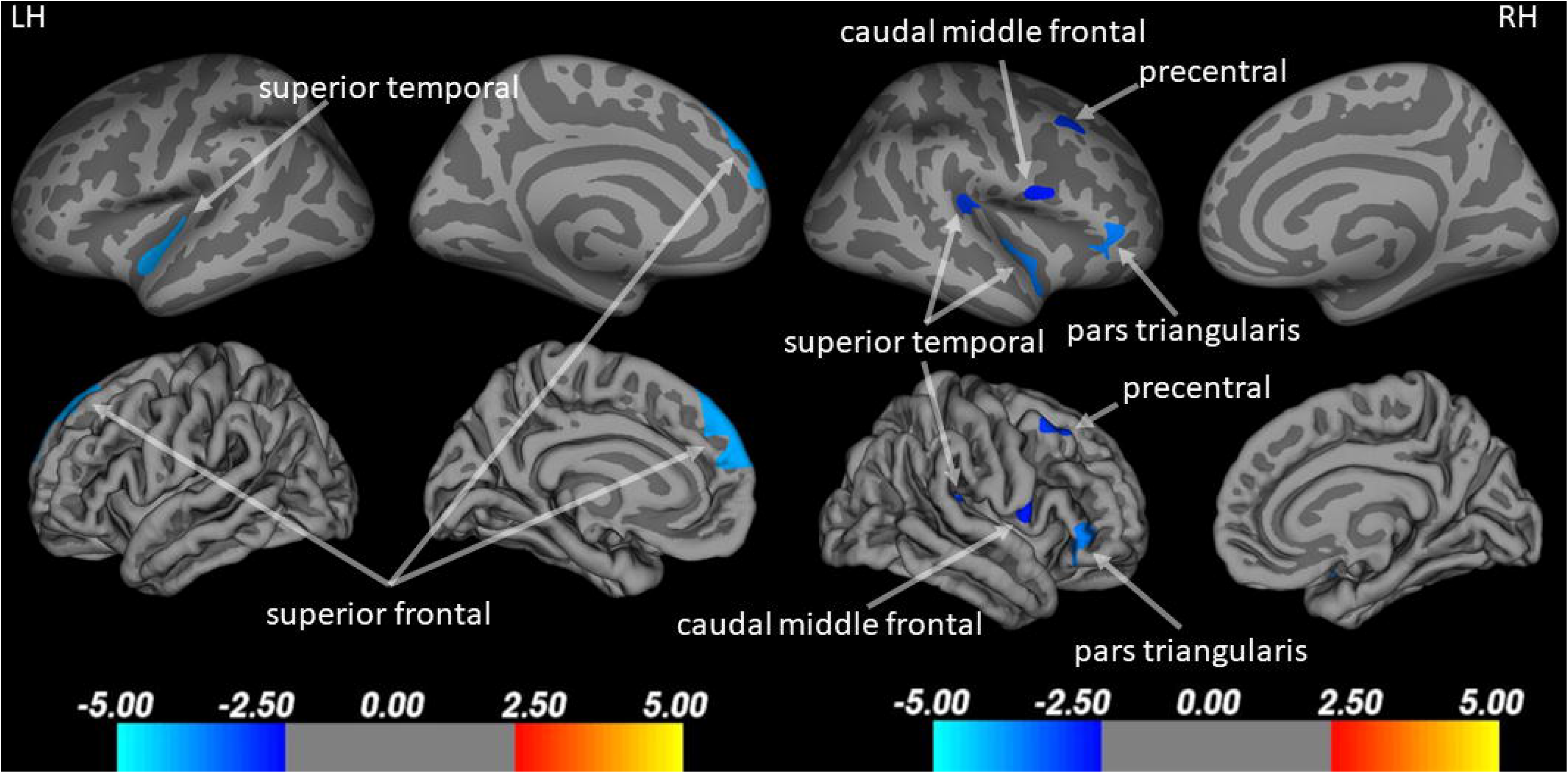
Results of the whole-brain FreeSurfer analysis for cortical thickness (CT) including the two age groups. Results are displayed on the inflated and the pial surface. Blue clusters refer to significant negative correlations (greater CT associated with better speech-in-noise perception). No significant positive correlations (red) were found. LH= left hemisphere, RH= right hemisphere.

Independent samples *t* tests revealed that CT was different between age groups in all regions extracted based on the significant results from the whole-brain analysis reported above (see Table 2).

**Table 2:** Age group differences in CT of regions of interest derived from the whole-brain analysis with speech-in-noise reception thresholds as predictors (*p*-values Bonferroni corrected). \**p* <.05, \*\**p* <.01, \*\*\**p*<.001.

|  | Older adults |  | Younger adults |  |  |  |
| --- | --- | --- | --- | --- | --- | --- |
|  | M | SD | M | SD | *t* | *p* |
| **LH** |  |  |  |  |  |  |
| superior frontal | 2.754 | 0.25 | 3.062 | 0.2 | t(36)=-3.885 | .003** |
| superior temporal | 2.309 | 0.23 | 2.655 | 0.19 | t(36)=-4.624 | <.001*** |
| **RH** |  |  |  |  |  |  |
| superior temporal | 2.493 | 0.24 | 2.852 | 0.22 | t(36)=-4.43 | .001** |
| pars triangularis | 2.471 | 0.22 | 2.727 | 0.16 | t(36)=-3.615 | .006** |
| transverse temporal | 2.122 | 0.21 | 2.451 | 0.19 | t(36)=-4.772 | <.001*** |
| caudal middle frontal | 2.608 | 0.24 | 2.882 | 0.27 | t(36)=-3.168 | .022* |
| precentral | 2.51 | 0.23 | 2.951 | 0.2 | t(36)=-5.917 | <.001*** |

Consistently, the OA’s cortex was thinner than those of the YA (Figure 4) in those brain regions suggesting that age-related cortical thinning was related to the lower SiN performance in OA compared to YA.

**Figure 4:**
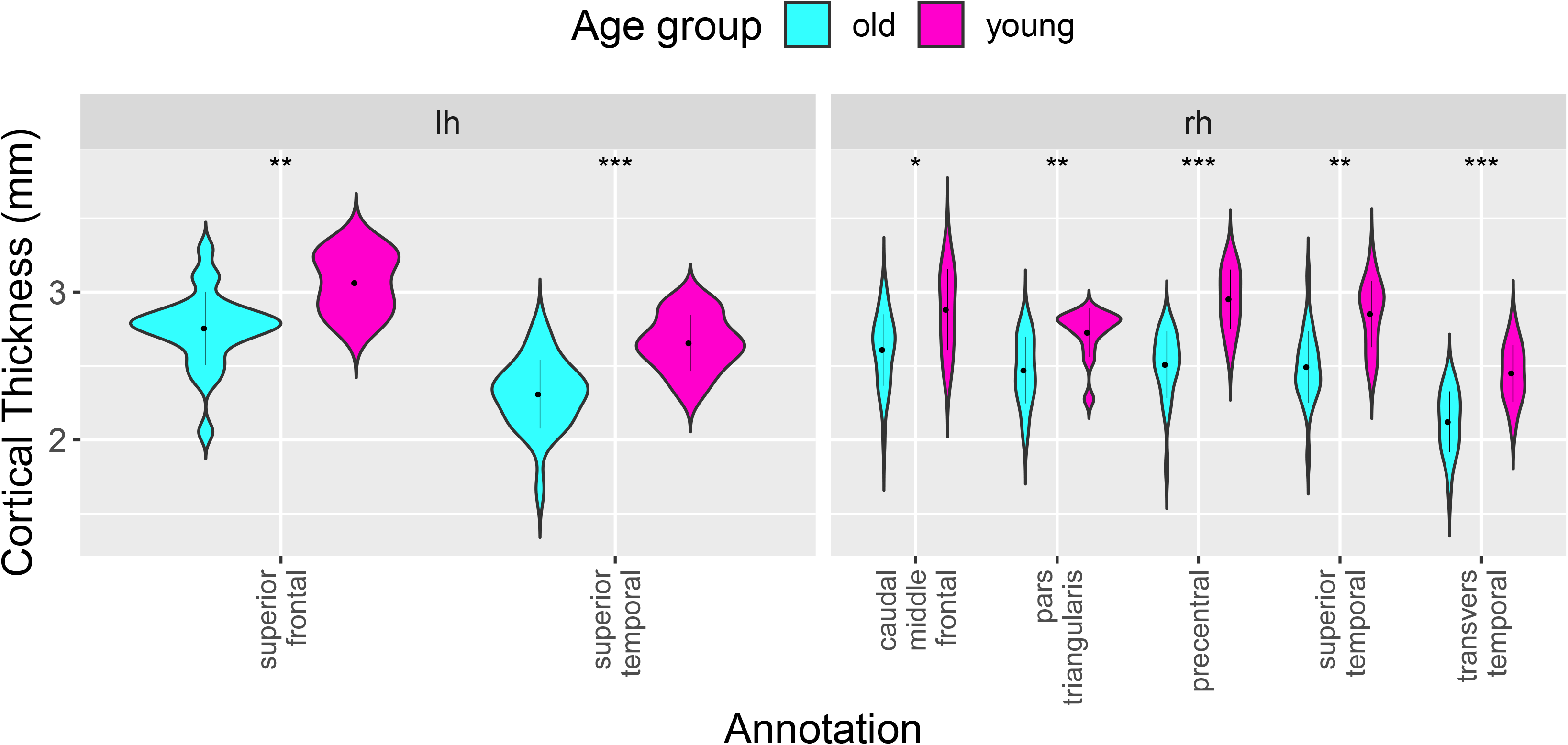
Age group differences in cortical thickness (CT) of brain regions associated with speech-in-noise perception (see Figure 3 and Table 1). lh= left hemisphere, rh= right hemisphere. \**p*<.05, \*\**p*<.01, \*\*\**p*<.001.

### 3.3. Thinner left superior frontal lobe associated with lower SiN recognition in older, but not younger adults

The regression model, testing to what degree age moderated the associations between CT in the above reported brain regions and SiN recognition (F(17, 20)=16.98, *p*<.001, adjusted R^2^=0.88), indicated that a thicker left superior frontal cortex predicted better SiN recognition (β=0.24, *p*=.041) in older adults only (see Figure 5). No age-moderating effects were found for the other brain regions reported in paragraph 3.2. These results suggest that greater age-related cortical thinning in the left superior frontal lobe is detrimental for SiN recognition in older adults.

**Figure 5:**
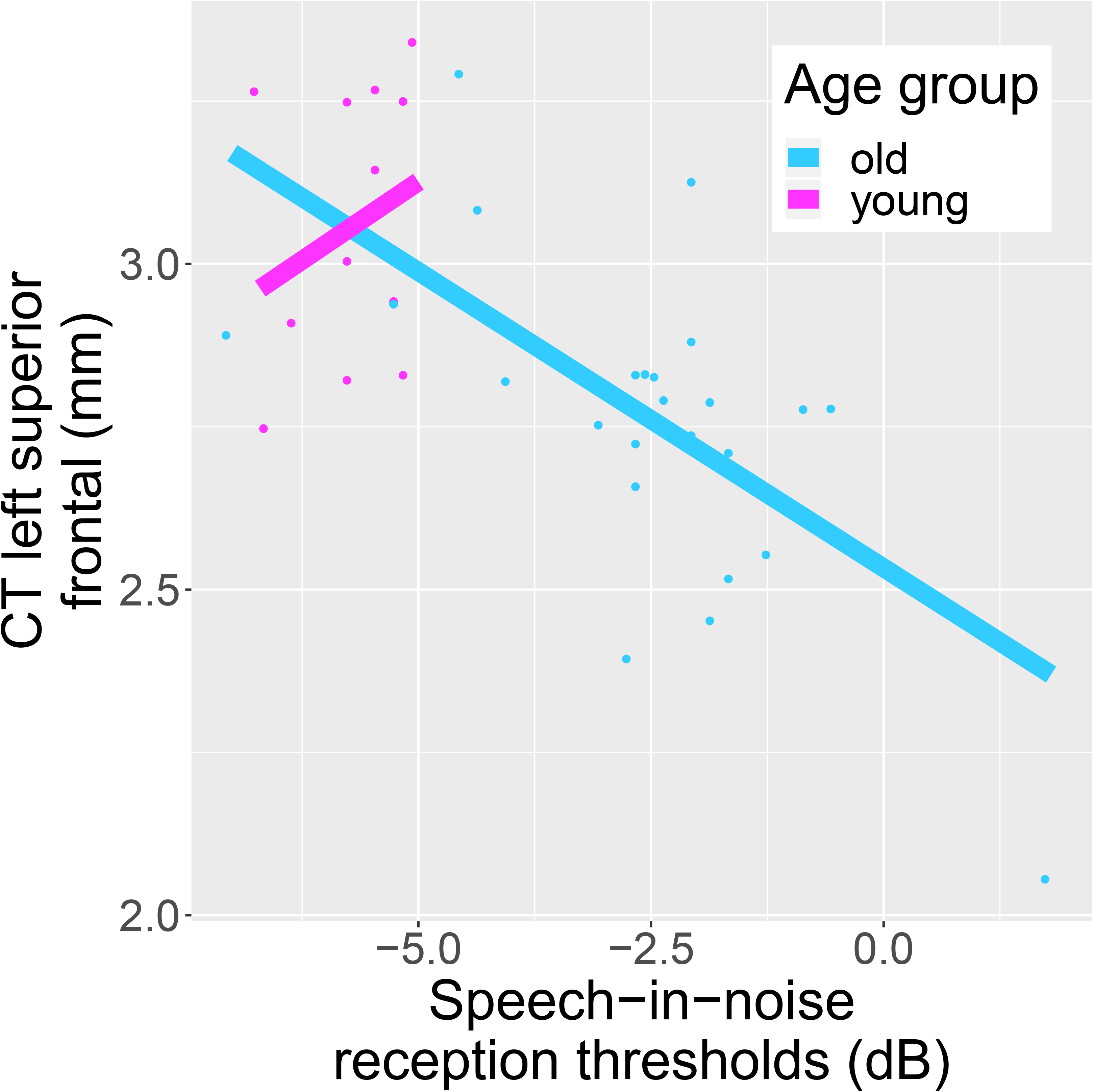
Age significantly moderates the association between cortical thickness (CT) and speech-in-noise (SiN) perception. In older adults (OA) only, those with greater CT in the left superior frontal lobe also perform better in the SiN perception task. This association was not significant in the younger age group.

An open question was to what extent greater pure-tone hearing loss (i.e., greater PTA) in the OA compared to the YA was related to the stronger involvement of the left superior frontal lobe in SiN recognition in older adults. The reported regression analysis in paragraph 3.3. was statistically controlled for PTA suggesting that greater pure-tone hearing loss in the OA did not significantly contribute to the findings. In order to further test whether individual differences in PTA within the OA group moderated the association between CT of the left superior frontal lobe and SiN recognition, we included PTA as a moderator in the regression analysis. The regression did not reach significance (F(3,21)=1.62, *p*=.22, adjusted R^2^=.07). Thus, we did not find evidence indicating that the association between CT in the left superior frontal lobe and SiN recognition occurred because of individual differences in pure-tone hearing loss.

### 3.4. Working memory moderates the association between CT in the left superior frontal lobe and SiN abilities in older adults

Further, we tested the hypothesis, that individual differences in working memory was moderating the association between CT in the left superior frontal lobe and SiN recognition in older adults. The regression analysis revealed that working memory (*β*=−0.41, *p*=.013) moderated the association between CT in the left superior frontal gyrus and SiN recognition in the OA (F(3,20)=8.66, *p*<.001, adjusted R^2^=0.5). There was no significant direct association between working memory capacity and SiN in the OA (*r*=.10, *p*=65). The association between a decrease in CT in the left superior frontal gyrus and greater SiN reception thresholds was stronger for individuals with higher working memory capacity within the older group (see Figure 6). Individual differences in working memory capacity, but not, might therefore drive the relevance of structural integrity of the left superior frontal gyrus in SiN recognition in older adults. Thus, our results suggest that older adults with higher working memory capacity might be better able to benefit from high structural integrity in the left superior frontal gyrus in order to perform well in a SiN recognition task.

**Figure 6:**
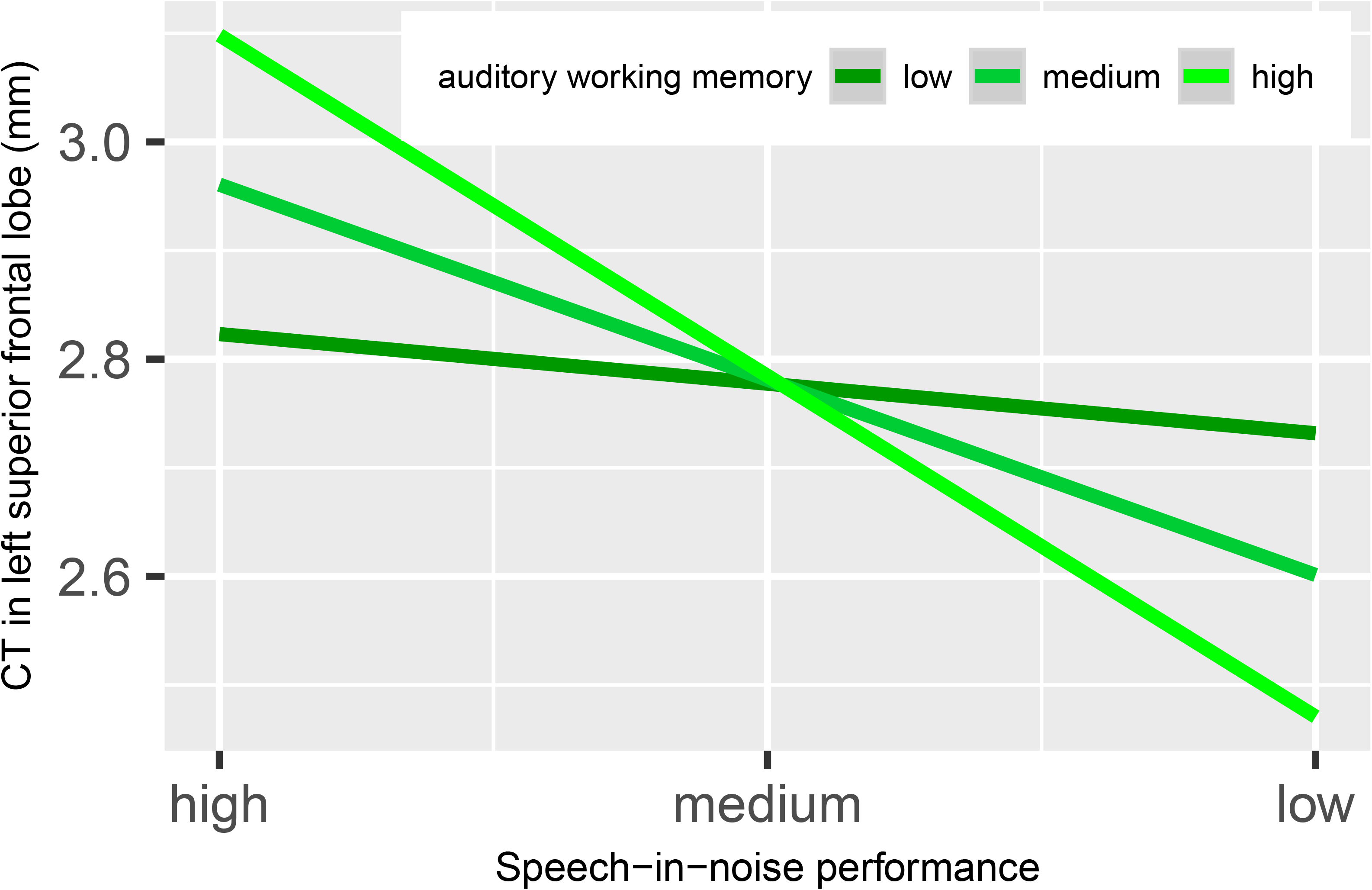
Working memory significantly moderates the association between cortical thickness (CT) in the left superior frontal lobe and speech-in-noise (SiN) perception in older adults. The relationship between SiN and CT in the left superior frontal lobe was stronger for older adults with higher working memory capacity compared to those with lower working memory abilities.

## 4. Discussion

Our work combined two separate lines of research showing that 1) age-related structural decline in auditory- and cognitive-related brain regions as well as 2) age-related cognitive decline are associated with the typically reported strong decline of SiN recognition in older adults. We consider it important to study the two factors in combination to find a suitable target for an intervention, because of the broad consequences of hearing and speech perception problems in older adults. Thus, the long-term goal of this research is to contribute to the development of feasible interventions targeting not only the ear (i.e., by hearing aids), but focusing also on neural and/or cognitive aspects of auditory processing difficulties in older adults.

### 4.1. Interacting effect of age-related cortical thinning and working memory decline in SiN recognition difficulties

In our sample, and in previous research (Dubno et al., 1984; Giroud et al., 2018; Gordon-Salant and Fitzgibbons, 1993; Killion and Niquette, 2000), there is a strong decline of SiN recognition in older compared to younger adults independently of pure-tone hearing loss. In our sample, the age group difference equals about 3.2 dB SNR on average. Our sample of older adults displayed consistent age-related atrophy in SiN-relevant regions such as the bilateral superior temporal regions and right inferior frontal and precentral regions. In other words, those who demonstrate greater thinning in those brain regions performed poorer in the SiN task. Our data further revealed the relevance of a structurally intact left superior frontal lobe for older adults’ SiN recognition because greater age-related thinning in the left superior frontal lobe was associated with lower SiN performance in older adults specifically. Importantly, this association between CT in the left superior frontal lobe and SiN recognition was moderated by individual working memory capacity. It seems that those older adults who have greater working memory capacity tend to benefit more from a structurally intact (i.e., less affected by age-related thinning) left superior frontal lobe to compensate for the greater challenge related to the processing of speech in background noise.

### 4.2. Frontal cortical thickness as a correlate for successful SiN recognition

Notably, all the neurostructural correlates of SiN recognition in this study were related to changes in cortical thickness and not cortical surface area. Younger adults were found to consistently have a thicker cortex. These findings corroborate previous observations in which lower cortical thickness and decreased cortical volume correlated with lower SiN recognition (i.e., higher SiN recognition thresholds) (Giroud et al., 2018; Rudner et al., 2019; Wong et al., 2010). This result speaks to the notion of CT and CSA as two independent anatomical traits (Meyer et al., 2016; Winkler et al., 2010). The interpretation is in line with a proposal of the Radial Unit Hypothesis describing different developmental trajectories of CT and CSA during prenatal brain maturation (Rakic, 1995, 1988). In the current study, changes in CT seem to reflect age-related processes of cortical atrophy, while CSA does not (Engvig et al., 2010; Giroud et al., 2018; Storsve et al., 2014). Our findings therefore underscore the possibility that the individual degree of age-related cortical thinning covaries with difficulties in SiN recognition in older adults.

Another explanation for the CT-SiN association stems from research with hearing-impaired older adults. In older and hard-of-hearing individuals, such variability in age-related neuroanatomical alterations might also occur as a consequence of reduced sensory input to the auditory pathways and eventually to the auditory cortex. In fact, longitudinal studies have demonstrated that greater pure-tone hearing loss was associated with greater gray matter volume loss (Lin et al., 2014; Xu et al., 2019) in auditory-related brain regions (e.g. bilateral STG) and areas associated with cognitive processing (e.g. the parahippocampus and the hippocampus) as well as lateral ventricle expansion (Eckert et al., 2019). Also cross-sectional work has shown that pure-tone hearing loss is related to neuroanatomical alterations of gray matter volume and thickness (Alfandari et al., 2018; Eckert et al., 2012; Rigters et al., 2018, 2017; Tuwaig et al., 2017; Rudner et al., 2019; Uchida et al., 2018; Ren et al., 2018; Armstrong et al., 2019; Neuschwander et al., 2019; Giroud et al., in prep., but see Profant et al., 2014). However, in our participants, only mild pure-tone hearing loss was evident making the first explanation for the CT-SiN correlations more likely. Future longitudinal studies in older adults with only mild or subclinical pure-tone hearing loss should be conducted to find out to what degree such small losses in pure-tone hearing could already lead to decline in brain structure across the lifespan.

Brain areas whose thicknesses were significantly correlated with SiN recognition were found in peri-sylvian and frontal regions, concurring with previous neuroanatomical findings (Giroud et al., 2018; Rudner et al., 2019; Wong et al., 2010). Besides the undisputed relevance of auditory superior temporal regions for SiN processing, CT in the right pars triangularis was correlated with SiN recognition across all participants. To our knowledge, the specific contribution of the right pars triangularis in speech processing is unclear to date. FMRI responses in the right IFG have previously been related to mental repair of spoken sentences (Meyer et al., 2000). Thus, it could be speculated that the right IFG is involved in the mental amplification of auditory input due to effortful filtering of the speech signal from the background noise. An alternative interpretation however could be that the right IFG supports attentional control and inhibition (Aron et al., 2004; Hampshire et al., 2010) pointing to the role of this region in executive functions when recognizing SiN.

### 4.3. The left superior frontal gyrus as a working memory allocator for SiN recognition in older adults

Importantly, cortical thinning of the left superior frontal gyrus seems to play an important role in the age-related decline of SiN recognition. Similar to our sample, this association, which was exclusively found in older participants irrespective of pure-tone thresholds, was reported previously (Wong et al., 2010). It was interpreted as reflecting a decline of cognition, particularly working memory, leading to a decline in older listener’s SiN recognition. Based on neurofunctional studies showing an involvement of the left superior frontal gyrus in working memory (Cornette et al., 2001; du Boisgueheneuc et al., 2006), these results overall suggest that fewer age-related neuromorphological anomalies in the left superior frontal gyrus might improve working memory-related processes during speech recognition in noise leading to better recognition performance. Even though Giroud et al. (2018) did not find this age-specific association between CT of the left superior frontal gyrus and SiN recognition, our moderation analysis supports this interpretation. Particularly individuals with high working memory capacity benefit from a neuroanatomically intact left superior frontal gyrus suggesting that this region may be responsible for the allocation of working memory resources during SiN processing. In other words, especially when working memory capacity is high enough to potentially facilitate SiN recognition in individuals with insufficient auditory structures, the intact left superior frontal gyrus will allocate those working memory resources to the task. Similarly, this interpretation would predict that when an older individual has low working memory capacity, the degree of anomaly in the left superior frontal gyrus does not matter to the same extent as in someone with high working memory capacity, because there are not enough working memory resources available to allocate to the task.

An alternative explanation is related to the inhibition hypothesis. The PFC is also involved in executive functions such as inhibitory control (Elliott, 2003), also of working memory contents (Hasher and Zacks, 1988). Thus, it is also possible that older adults with high working memory capacity keep more irrelevant information in mind (e.g., competing words during lexical access) leading to a higher need to inhibit incorrect words (Wong et al., 2010) which can be facilitated by a structurally intact left superior frontal lobe.

Notably though, in our sample working memory capacity did not directly correlate with SiN recognition (only by moderating the association with CT of the left superior lobe). This is in line with a previous study assessing to what extent working memory training would transfer to SiN recognition in older adults which did not find any significant improvement in SiN performance after training (Wayne et al., 2016). Further, it has been noted that in normal hearing younger listeners there is only weak evidence for the association between working memory and speech-in-noise recognition (Füllgrabe and Rosen, 2016a, 2016b). Also, the association depends strongly on the test environment, such as the working memory task modality and the speech in noise masker type (Besser et al., 2013). Finally, it has been reported that the working memory benefit stems mainly from the capitalization of contextual cues (Gordon-Salant and Cole, 2016), which were not present in our study. Thus, it has to be noted that the interaction between working memory and SiN recognition appears not to be always straight forward. In terms of finding a target for an intervention to improve speech processing in background noise for older adults, a working memory training might not be the primary target. Our study suggests that a more promising candidate could be a brain stimulation protocol improving frontal compensatory processes during SiN perception directly.

## Acknowledgments

This study was funded by the Swiss National Science Foundation (SNF, no. 105314_152905; no. 105319_169964 to MM) and by a postdoctoral “Forschungskredit” from the University of Zurich (no. FK-19-072 to NG). During the work on his dissertation, MK was a pre-doctoral fellow of LIFE (International Max Planck Research School on the Life Course; participating institutions: MPI for Human Development, Humboldt-Universität zu Berlin, Freie Universität Berlin, University of Michigan, University of Virginia, University of Zurich). Financial support by the Jacobs Foundation helped to conduct this research. Furthermore, this research was supported by the University Research Priority Program (URPP) ‘Dynamics of Healthy Aging’.

